# FedDP: Secure Federated Learning for Disease Prediction with Imbalanced Genetic Data

**DOI:** 10.1101/2023.01.17.524409

**Authors:** Bin Li, Hongchang Gao, Xinghua Shi

## Abstract

It is challenging to share and aggregate biomedical data distributed among multiple institutions or computing resources due to various concerns including data privacy, security, and confidentiality. The federated Learning (FL) schema can effectively enable multiple institutions jointly perform machine learning by training a robust model with local data to satisfy the requirement of user privacy protection as well as data security. However, conventional FL methods are exposed to the risk of gradient leakage and cannot be directly applied to genetic data since they cannot address the unique challenges of data imbalance typically seen in genomics. To provide secure and efficient disease prediction based on genetic data distributed across multiple parties, we propose an FL framework enhanced with differential privacy (FedDP) on trained model parameters. In FedDP, local models can be trained among multiple local-hold genetic data with efficient secure and privacy-preserving techniques. The key idea of FedDP is to deploy differential privacy on compressed intermediate gradients that are computed and transmitted by optimizers from local parties. In addition, the unique weighted minmax loss in FedDP is able to address the difficulties of prediction for highly imbalanced genetic datasets. Our experiments on multiple genetic datasets demonstrate that FedDP provides a powerful tool to implement and evaluate various strategies in support of privacy preservation and model performance guarantee to overcome data imbalance.

## 1 Introduction

Cancer patients may get their genome sequenced to help determine the subtypes of their tumor based on the genotypes or molecular markers to receive personalized therapy and care. An artificial intelligence (AI) algorithm can be used to take in genome, and calculate a risk score, classify her tumor into particular subtypes, determine treatment strategies, and predict clinical outcomes[30]. To provide disease prediction aforementioned in precision medicine, sharable large-scale data resources that include a large collection of patient data from diverse populations, geographical and ethical groups would be desirable[20]. However, such data are usually distributed across multiple institutions, health providers or computing infrastructure which makes such data not easily shareable due to various constraints including privacy concerns[25][21]. Hence, it is challenging and sometimes infeasible to aggregate and share a large amount of data that is required and necessary to develop robust, secure, and trustworthy AI systems for biomedical research and clinical practices[12].

With the recent advancement of AI and machine learning (ML), the identification of diseases and the diagnosis of ailments are at the forefront of the applications of AI/ML in the medical field[7]. In high-demand areas such as cancer identification and treatment, it is a research frontier in leveraging AI/ML to design more effective therapies based on the combination of individual genetic data with health information, empowered by predictive analysis[8]. According to a 2015 report by Pharmaceutical Research and Manufacturers of America([2]), more than 800 drugs and vaccines for the treatment of cancer are in clinical trials.

However, the effectiveness of ML algorithms often depends on the amount of data that is used to train these algorithms. A large dataset of high quality will help train a robust ML algorithm to yield promising results. Therefore, the present core issue at the intersection of ML and medicine is to safely collect and aggregate lots of different types of data for better analysis, prevention, and treatment of individuals. In many cases, data owners could be reluctant to share their data (e.g., genomic data from patients) due to constraints about data privacy and patient confidentiality, together with restrictions of privacy policies and patient consents[9]. To provide secure and privacy-preserving protection of data and analytics, there have been various methods developed including differential privacy (DP) and cryptographic solutions in health domains. As a classical privacy-preserving method, DP have been applied to preserve genome privacy in genome-wide association studies (GWAS)[22]. For example, [14] developed privacy-preserving algorithms for computing the number and location of single nucleotide polymorphisms (SNPs) that are significantly associated with diseases. [26] proposed a method that allows for the release of aggregated GWAS data without compromising an individual’s privacy. Various DP mechanisms have been developed [17] to preserve the privacy of various ML algorithms, including logistic regression, random forest [18] and deep neural networks [23]. Moreover, [3] developed a differentially private stochastic gradient descent (SGD) algorithm for the TensorFlow framework.

Federated Learning (FL) can effectively help multiple institutions to perform data usage and ML modeling under the requirements of user privacy protection, data security, and government regulations[27][10]. The model based on both FL and DP has significant privacy protection capabilities in the direction of medical diagnosis[19][4]. The main idea is to share model parameters with DP[31], not original data, aggregating them regularly after local training steps, that will eventually lead to a shared global model with secure training on data distributed at multiple parties[23]. However, genomic data are highly imbalanced and typically distributed in different institutions, limiting the possibility of training well-performing FL models to benefit healthcare decision-making[11][13]. Specifically, those FL methods typically optimize the accuracy-induced objective function, e.g., the cross-entropy loss function, whose incapability of handling imbalanced data has been confirmed by numerous studies [16][6]. Hence, it is necessary to develop new FL approaches to address the issue of imbalanced data distribution as demonstrated in genomic analysis[5].

In this study, we propose a federated learning framework incorporated with differential privacy (FedDP) to provide differentially private federated learning for disease prediction, with optimization strategies deployed to handle imbalanced data. In FedDP, differential privacy was deployed on model parameters during the transmission instead of local datasets. Meantime, we used a weighted min-max optimizer to give assistance to enhance the capacity of secure federated learning on imbalanced data which is commonly found in genomic and biomedical datasets. Our empirical results on various datasets have shown that FedDP is efficient in support of privacy-preserving disease prediction, even when data are imbalanced among different parties.

The contribution of this study is summarized below.

**First**, we investigate the shortcomings of current federated models and develop a strategy to strike a trade-off between the security and performance of federated models. Instead of imposing privacy preservation on the original dataset, DP noises are attached to model gradients in each federated transmission from local parties to the global server. Our FedDP framework is a privacy-preserving FL model that shares model gradients with (*ϵ, δ*)-DP instead of the original data, aggregating them regularly after local training steps, thus leading to a shared global model. Results show that FedDP can be deployed under a strict privacy budget with little sacrifice of model accuracy.

**Second**, The proposed FedDp employs the AUC-induced min-max loss function rather than optimizing the accuracy-induced loss (e.g., cross-entropy), which can address the issue of imbalanced data distribution in genetic data.

**Third**, we demonstrate the proposed FedDP by applying it to two breast cancer datasets for cancer prediction. We evaluate the performance of the FedDP framework in two scenarios where different parties have balanced datasets and imbalanced datasets respectively. Experimental results show that FedDP performs well under the strong privacy budget in both two cases, indicating the generality and effectiveness of the FedDP framework for disease prediction on genetic datasets.

## 2 FedDP Framework

In this section, we present our privacy-preserving federated learning framework that combines FL with DP (FedDP) for disease prediction.

### 2.1 Federated Learning Scheme

Federated Learning (FL) is a promising learning paradigm for preserving privacy when learning on distributed data. In FL, rather than sharing the raw data, a machine learning model is shared among participants in a privacy-preserving way. The private information in local data can be preserved, and the machine learning model can achieve good generalization performance under this schema. To demonstrate the idea, the federated framework proposed in this paper includes one centralized server and multiple parties.

As simplified in **Figure 1**, we illustrate a simplistic setting where this exemplar FL system is composed of one centralized server and three parties deployed at different devices.

**Fig. 1.**
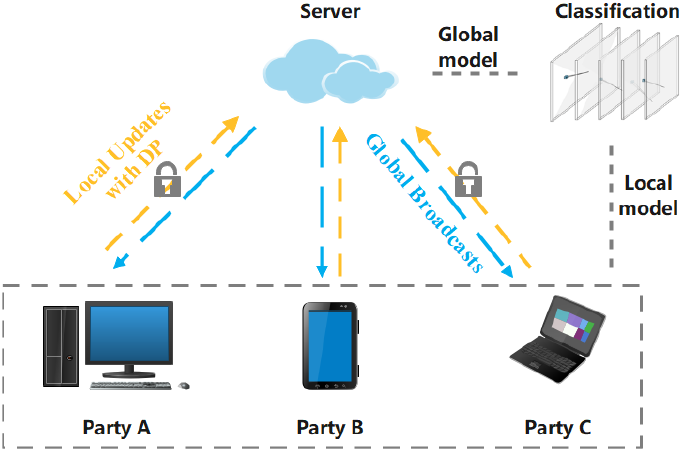
An illustration of a federated framework. This FL framework includes a central trusted server and three collaborating parties (A,B,C) that can be deployed on different computing devices.

Each federated round (as shown in **Figure 1**) is conducted following four steps:

1. Initialize the parameters and data of each party and the server.

2. Each party trains the model separately based on their local data, and then sends these intermediate parameters to the server.

3. The global optimizer applies an averaged update to the global model at the server end.

4. The central server broadcasts the aggregated parameters to the parties at the end of each federated round. Afterward, each party updates its local models with these new parameters.

#### Structure of FedDP

Based on this centralized FL framework, we employed the DP mechanism for securing the intermediate parameters in the transmission process. In addition, weights from each local DL network will not be transmitted in a federated process, instead, gradients computed by local optimizers are selected as intermediate parameters with (*ϵ, δ*) DP noises. This framework which combines gradients-transmitted FL and DP is called FedDP. The overall framework of FedDP is shown in **Figure 2**.

**Fig. 2.**
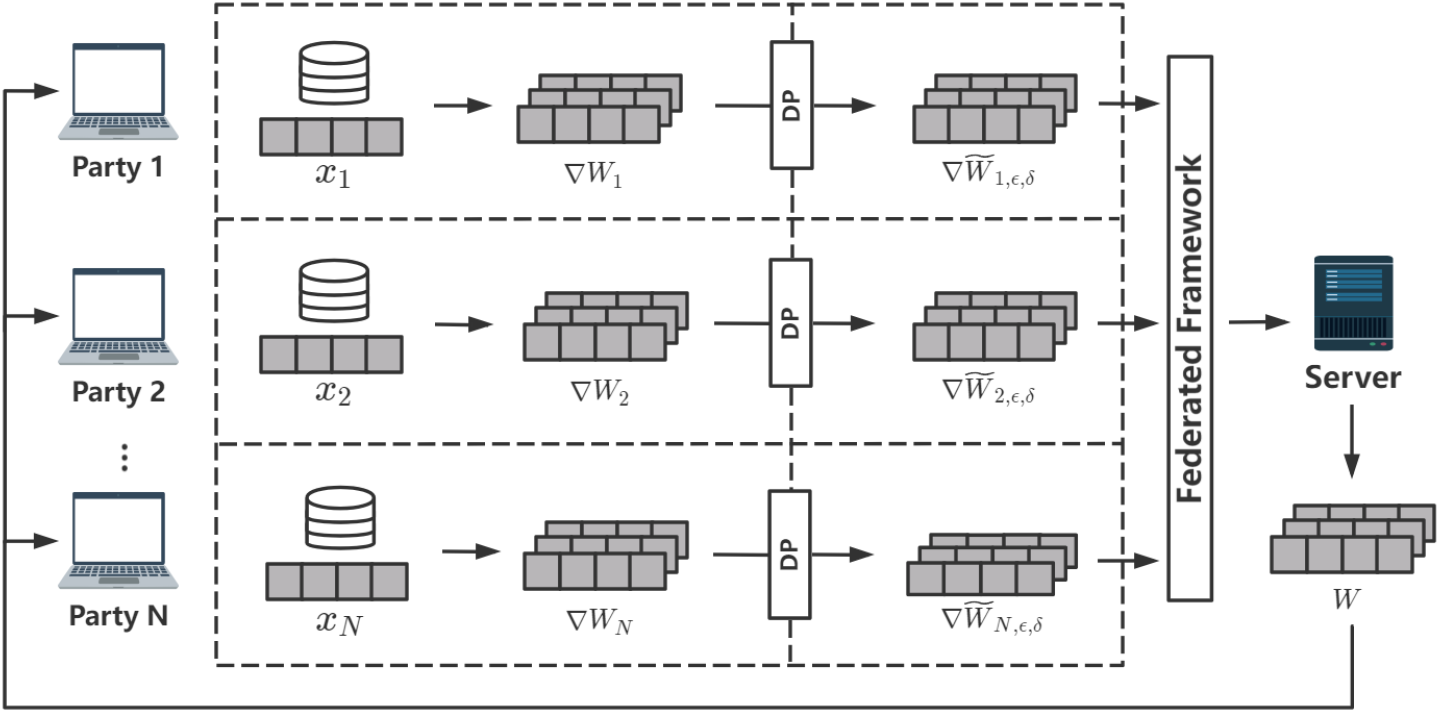
An overall framework of FedDP. Local data *x*_*i*_ distributed at each party is independent of each other First, local gradients ∇*W*_*i*_ are generated in parallel by each local optimizer deployed at each party and trained on each local dataset. Then, the server gathers all gradients 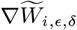 from multiple parties after (*ϵ, δ*) DP was applied to original gradients ∇*W*_*i*_. Finally, global weights are computed and broadcast to each party at the end of each round by aggregating all 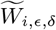.

It should be noted that there are two kinds of optimizers in FedDP: local optimizers with standard (*ϵ, δ*)-DP and a server optimizer. The local optimizer is only used to compute local model updates at each party. We add DP to the parameters to be transmitted from local optimizers to ensure the ability of privacy-preserving. In this perspective, this distributed architecture ensures that each party does not need to share their data with others..

#### Training FedDP

For *N* parties, local data *x*_*i*_ were held on their own side. Thus, the privacy of original data was secured by global invisibility. Each local model computes gradients of weights in every layer. Gaussian DP mechanism is applied to intermediate gradients to escalate the privacy preservation under the potential attacks. The central server received all the gradients 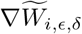 from each party. The global weights *W* are computed by the global model by aggregating local gradients 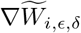 Subsequently, local updating occurred once local parties received the global weights broadcast by the central server.

### 2.2 Aggregation Method

For federated model training, gradients computed from local datasets are communicated to the central server at the end of each round. The pseudo-code of the proposed differentially private federated training procedure is depicted in Algorithm 1 (see Appendices). In FedDP, global gradients were aggregated by averaging all gradients learned from local parties. Multiple parties could set different aggregating weights regarding their relative data sizes.

The proposed procedure protects the per-sample privacy of each participating party dataset. DP is added by parties individually. In this way, the dataset sizes can be very different among the parties. Dependent on the setting of the sampling strategy, the sampling ratio can be different from local to local. Thus the privacy loss may vary on each local dataset. For local updating, each local party should keep a record of the spent privacy along each update, and stops its update to the central server once the predefined privacy threshold *E* is reached.

## 3 Data and Experiments

In this section, we present our experimental results. First, we illustrate the details of three datasets we used in this research. Then we introduce the optimization method and evaluation metrics of FedDP. Lastly, we show the performance and hyperparameter settings of FedDP in two cases that performance with balanced data distribution and imbalanced data distribution for each local party. The whole process of FL is simulated on one high-performance clustering server that has two NVIDIA Tesla V100 GPUs. We evaluate the performance of the proposed algorithm under a varying (*ϵ,δ*)-DP privacy budget. The grid-search method is performed to fine-tune the parameters in experiments. The result under each set of parameters was averaged 5 times.

### 3.1 Datasets

To investigate the contribution of privacy preservation for a federated learning framework, we evaluated FedDP with two different models on two real-world cancer datasets.

The Cancer Genome Atlas (TCGA), which is a project overseen by the National Cancer Institute and the National Human Genome Research Institute to apply high-throughput genome analysis techniques to help people have a better understanding of cancer as well as improve the ability to prevent, diagnose and treat cancer. To evaluate FedDP for the cancer prediction task, we utilized two TCGA datasets, BC-TCGA and TCGA-BRCA. Both datasets label the samples as two categories, that is, positive (breast-tumor) samples and negative (normal) samples. The BC-TCGA dataset is a single dataset containing 61 normal samples and 529 breast-tumor samples. Each sample has gene expression profiles 17,814 genes. While another TCGA-BRCA dataset, has a larger sample size including 1217 samples (1103 breast-tumor samples, 114 normal samples) and each sample has expression quantification of 60,843 genes. Thus, these two TCGA datasets enable us to test FedDP on a smaller dataset and a larger dataset to explore the potential performance of FedDP for cancer prediction.

We randomly split datasets into three sub-datasets. Each sub-dataset is divided into one training set and one test set using label stratification. We filled missing data with zeros to construct the input datasets and all data are normalized into the range of zero to one [24]. Furthermore, to reduce the feature space for each dataset, we sorted features by fisher score to find a subset of features. Feature with a higher score means the distances between data points on this feature in different classes are as large as possible, while the distances between data points in the same class are as small as possible. Moreover, we further extracted 2000 sorted features with the least correlated coefficient with others since the fisher score ignores the correlation between each feature. Figure 1 in the supplementary material provides more details of feature selection in this experiment.

### 3.2 Model Architecture

In order to predict cancer types based on the genetic dataset, we built a two-layered CNN model for local training (as shown in **Figure 3**). Since the input data are a two-dimensional matrix, we constructed two Cov1D layers in which the size is 32× 2000× 16. We observed that the classification task is prone to overfitting because of the large number of input features. Thus, we reduced the number of effective weights by deploying L1 regularization at the end of each convolution layer. In addition, we constructed the logistic regression model with L1 regularization (lasso) as a comparison.

**Fig. 3.**
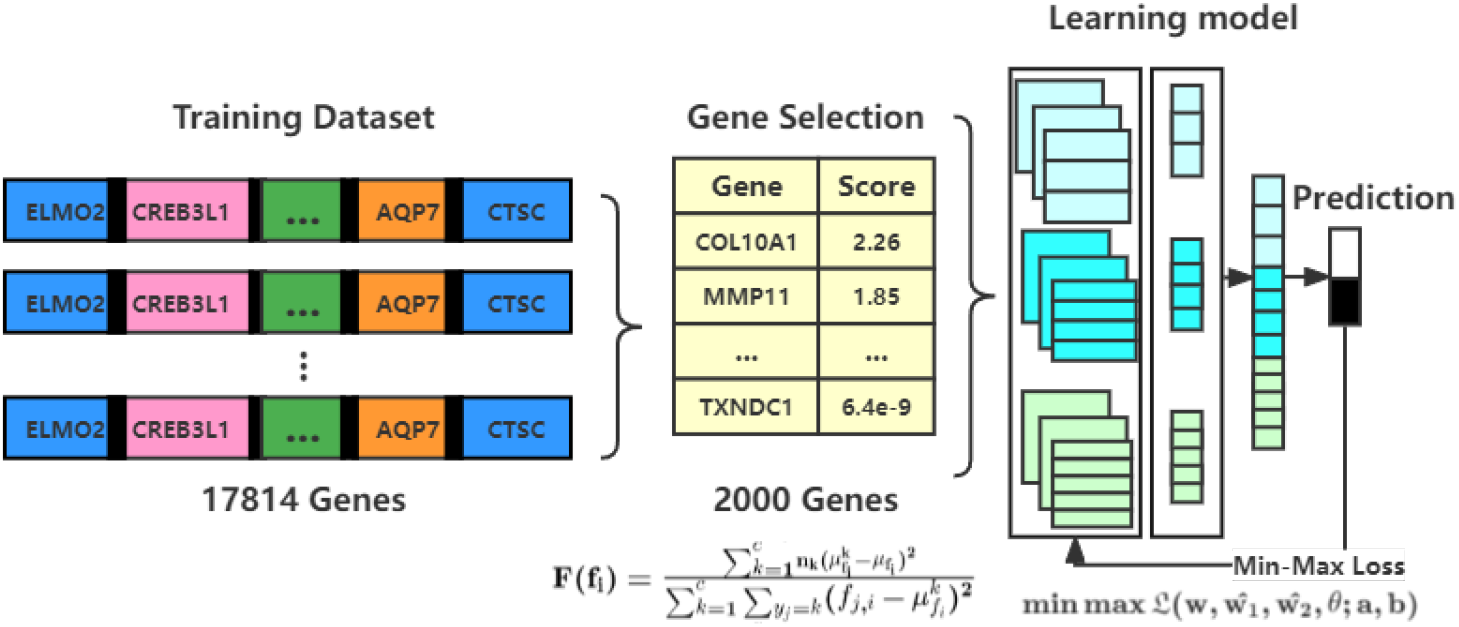
Illustration of model structure including feature scoring and selection. The CNN model is composed of two convolutional layers with two pooling layers and two dense layers. L1 regularization is applied to each layer to reduce the number of effective weights. Weighted min-max loss is employed to address label imbalance against overfitting.

### 3.3 Optimization Weighted Min-Max Loss

The conventional accuracy-optimized cross-entropy loss function is indeed applicable to binary and multi-class classification problems. However, it is challenging to learn a well-performing classifier with imbalanced genetic data. The model is prone to run into a dead-end in which all test samples are predicted as the major category, in this way, the loss function no longer drops because the accuracy is very high (very low precision or recall) at a certain point. Therefore, recent studies were focused on optimizing the area under the receiver operator curve (AUC) which is a reasonable metric for imbalanced-based-data classification tasks. The main feature of AUC is that it aggregates across different threshold values for binary prediction, separating the issues of threshold setting from predictive power. Meanwhile, AUC considers both precision and recall metrics to learn a comprehensive model across imbalanced data. Thus, it is rational to directly optimize the AUC score, rather than the cross-entropy loss function. Specifically, [28] developed the following minimax loss function for the AUC maximization problem:

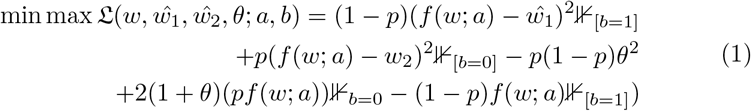

where *ŵ* ∈ ℝ ^*d*^ denotes the weights of model *f, ŵ*_1_ ∈ ℝ, *ŵ*_2_ ∈ ℝ, *θ* ∈ ℝ are additional parameters to compute AUC score, *a* is the feature of samples and *b* represents the label. *p* is the rate of positive samples to all samples and ⊮ is an indicator function.

#### Optimizer

The mini-batch ADAM SGD is used as local optimizers at local ends. On the one hand, the SGD optimizer can significantly reduce the time to compute gradients at each state as well as has low chances to overfit. On the other hand, ADAM has the advantage that its learning rate is adaptively adjusted, which helps each party to learn a different learning rate based on its own data distribution, especially in cases where each party holds a highly diverse dataset.

### 3.4 Evaluation Metrics

Our solutions are based on conventional (*ϵ,δ*)-DP model implemented by the Tensorflow privacy framework[1]. The term *ϵ*, which is called privacy budget, is the metric to determine if a system has a strict privacy-preserving ability under DP. The *ϵ* calculated in Tensorflow privacy is influenced by factors below:

**–** Sampling rate. The sampling rate is represented as the ratio of Batch size to Sample size. Theoretically, a lower sampling rate always generates lower epsilon if other factors are constant, which means better privacy-preserving capabilities.

**–** Noise multiplier: Noise multiplier is the ratio of the standard deviation of the Gaussian noise to the l2-sensitivity of the function to which it is added. This governs the amount of noise added during training. Generally, more noise results in better privacy and lower utility. This generally has to be at least 0.3 to obtain rigorous privacy guarantees, but smaller values may still be acceptable for practical purposes.

**–** Delta(*δ*): Delta bounds the probability of an arbitrary change in model behavior. We can usually set this to a very small number (1e-7 or so) without compromising utility. A rule of thumb is to set it to be less than the inverse of the training data size. (We set delta to 1e-5 this time)

Several optimal parameter settings for the two models on two different datasets are shown in **Table 1**. Our experimental results show that the smaller the batch size is and the greater noise_multiplier is, the smaller the privacy budget is, which means the higher degree of privacy-preserving capability. Moreover, our results demonstrate that increasing batch sizes will lead to higher accuracy but low privacy security. Thus, the accuracy of disease prediction in FedDP is positively related to the noise multiplier and *δ*. Consequently, reducing batch sizes means that more time is consumed during training. Thus, it is vital to make a trade-off between privacy budget and accuracy which are chosen as evaluation metrics throughout the whole parameter tuning process.

**Table 1.** Performance of FedDP on BC-TCGA (*D*_1_) and TCGA-BRCA (*D*_2_). Each row represents the accuracy under certain hyperparameter settings. There are two kinds of models deployed in FedDP: logistic regression and two-layered CNN. The trade-off taken between accuracy and privacy budget epsilon should be taken to obtain the best performance.

| Noise Multiplier | Delta | Batch Size | Privacy Budget $\epsilon_{d1}$ | Privacy Budget $\epsilon_{d2}$ | Acc | | | |
| --- | --- | --- | --- | --- | --- | --- | --- | --- |
| | | | | | $D_1, LR$ | $D_2, LR$ | $D_1, CNN$ | $D_2, CNN$ |
| **1.5** | **1e-3** | **32** | **0.84** | **1.91** | **0.95** | **0.53** | **0.98** | **0.93** |
| 1.3 | 1e-3 | 32 | 1.11 | 2.35 | 0.96 | 0.62 | 0.98 | 0.93 |
| 1.1 | 1e-3 | 64 | 2.39 | 4.15 | 0.98 | 0.70 | 0.99 | 0.94 |
| 1.1 | 1e-3 | 32 | 1.6 | 3.11 | 0.98 | 0.75 | 0.99 | 0.93 |
| **1.0** | **1e-3** | **32** | **1.94** | **3.69** | **0.99** | **0.89** | **0.99** | **0.93** |
| 1.0 | 1e-3 | 32 | 3.11 | 4.89 | 0.99 | 0.92 | 0.99 | 0.96 |
| 0 | 0 | 32 | None | None | 0.99 | 0.92 | 0.99 | 0.98 |

### 3.5 Experimental Results

#### Prediction vs. Privacy

To explore how the DP impacts the accuracy of the model, we tuned 3 hyperparameters to find the best setting for *ϵ* by conducting a grid search. The results indicate the model performance deteriorates with the decrease of the privacy budget *ϵ*. Specifically, for the classification of the BC-TCGA dataset based on the logistic regression model (LR), to maintain higher accuracy rather than privacy, the learning rate of DP-SGD on the server is appropriate at 0.003, and the number of epochs per round on each party is 15, and the noise multiplier is set to 1.0. Based on this hyperparameter setting, we have achieved a 1.0 accuracy under the DP criteria that *ϵ* is 5.38 and *δ* is 1e-3. Moreover, if we increased the batch size for training dataset. There would be a higher privacy budget after calculation, which means all procedures will be secured under a much strict privacy criteria. Certainly, the performance reached max if there is no-level DP added to the transmission. However, this will be exposed to the risk of privacy leakage. On the contrary, the best privacy budget *ϵ* we obtained is 0.84, which means the whole training process is under very strict privacy preservation. This also sacrifices the accuracy in test processing, which has 0.98 accuracies. In fact, this result is relatively promising due to the complexity of the TC-BCGA dataset. For deep learning models, it is simple to find the classification boundary for the BC-TCGA dataset. For the TCGA-BRCA dataset, the LR model was not able to reach an accuracy greater than 0.6. That means finding a boundary for the TCGA-BRCA dataset is difficult. Our solution is a nonlinear CNN model on the TCGA-BRCA dataset. As the results shown in **Table 1** and Figure 4, within the best setting, we finally could achieve a 1.0 accuracy under the DP criteria that *ϵ* is 5.38 and *δ* is 1e-3. Furthermore, we can gain an even stricter privacy budget with 0.74 accuracies. In conclusion, the CNN with min-max loss has a better capability to protect privacy compared with the linear logistic regression model.

**Fig. 4.**
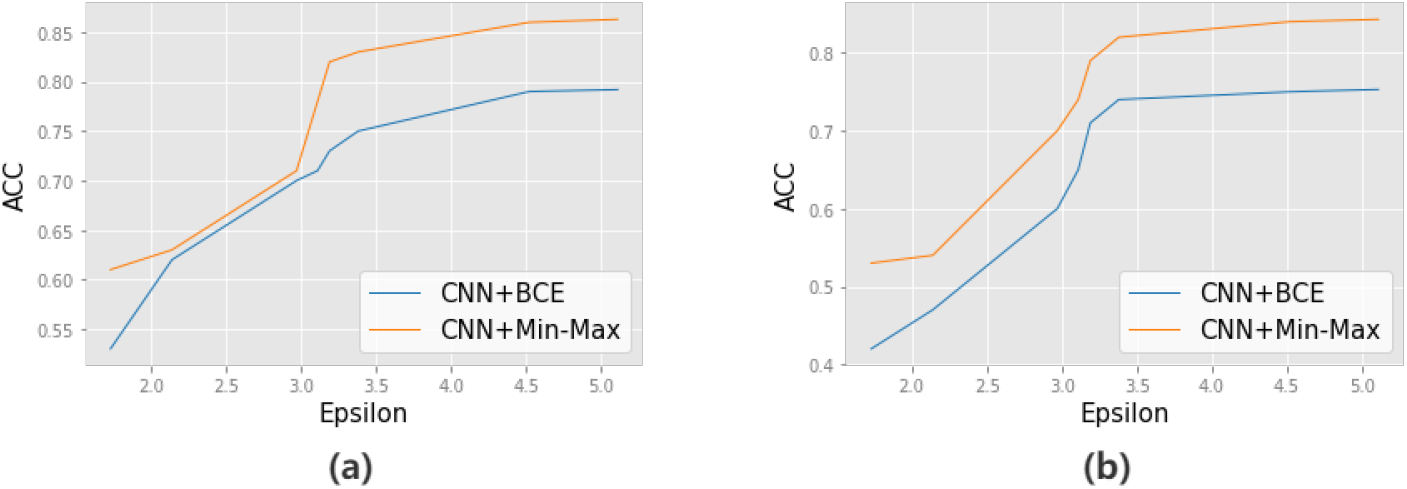
Model testing accuracy with different privacy budget epsilon values under the data-center setting with three parties: (a) Results on BC-TCGA dataset. (b) Results on TCGA-BRCA datasets.

#### Predicion for pathological stages

To explore the ability of FedDP to address multi-class prediction problems. We used another TCGA dataset containing gene expression values of kidney cancer for 1107 samples. Each sample has 60487 genes. The expression values correspond to RNA sequences in FPKM UQ formation. In addition, we integrated pathological stages for all cancer types into early, mid, and advanced stages based on the official categorical standards.

#### Addressing Label Imbalance

Since the dataset on each party is imbalanced, (i.e., 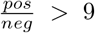) we can further explore if all parties collaboratively optimize the loss function to learn the best model parameter against imbalance. **Figure5** (a) shows the classification performance (AUC) of min-max loss compared with conventional binary cross-entropy (BCE) loss on BC-TCGA and TCGA-BRCA datasets. Furthermore, we evaluate FedDP with its precision and recall scores. **Figure 5** (b) shows that the precision is low and cannot gain much learning during the training process, which means BCE loss cannot address the imbalance problem because precision represents how accurate the prediction in the positive sample is. Low precision indicates there is a large amount of false positive predictions, the model is prone to predict all samples as positive. Accordingly, the AUC score of the model with BCE loss is around 0.5 meaning the model tends to be a “random prediction” model since it didn’t learn any relation between features and labels due to the high-imbalanced data. **Figure 5** (c) shows weighted min-max could overcome the data-imbalanced problem of two TCGA datasets, the precision score is improved fast during the training process as well. The shaded area reflects the fluctuation of learning among all parties since each local subdataset has a different data-imbalanced ratio.

**Fig. 5.**
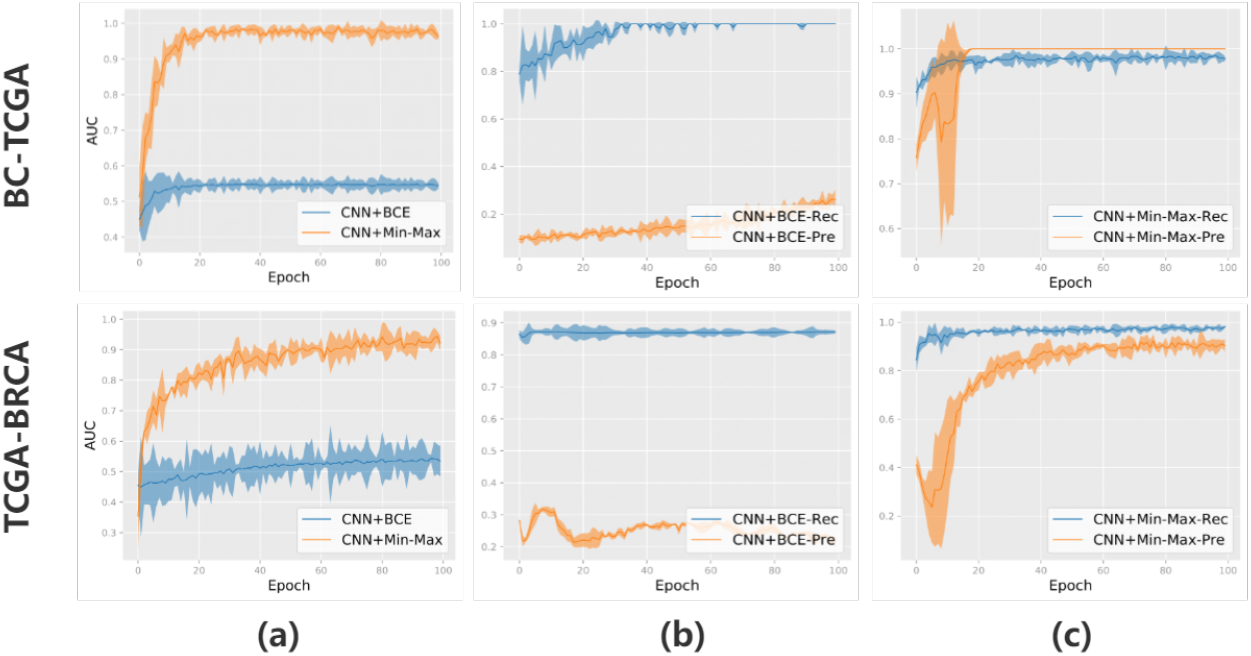
Improved cancer prediction results of the min-max optimization of CNN models under the data-center setting with three parties: (a) AUC scores of the min-max optimization (Orange) and a regular binary cross-entropy (BCE) optimization (Blue) on the same CNN model over 100 epochs. Light-shaded areas reflect the dispersion of AUC scores for the three parties. (b) The precision (orange) and recall (blue) score of the CNN model with BCE loss over 100 epochs. (c) Precision (orange) and recall (blue) score of the same CNN model with weighted min-max loss function over 100 epochs.

## 4 Conclusion and Future Work

In this paper, we propose a framework, FedDP, a privacy-preserving FL framework incorporating DP strategies. Instead of adding noises to the weights of the model at each federated round, noises are added to the gradients of each local model. We thus provide a more secure version of FL with DP, since gradients transmitted with DP are more secure than weights transmitted with DP. The CNN with DP-SGD achieved a good trade-off between accuracy and privacy budget as well as the consideration of time consumption. The weighted crossentropy loss has significantly improved the low accuracy problem caused by the label imbalance from genetic datasets. We tested our FedDP with imbalanced data load for each party. The model works well for a stringent privacy budget and the accuracy finally converged.

Although FedDP is a robust and flexible framework, it requires relatively high bandwidth for communications between the server and each party. The throughput capacity of central server is the bottleneck for such a FL framework. Furthermore, complicated deep models need more weights as well as gradients to be transferred in the FedDP framework.

Additionally, we can adjust the current FedDP architecture by removing the central server and extend it to a distributed network setting without a central server. One potential solution is that each party could share its encrypted gradients with its neighbors during the transmission so that the whole process can be implemented asynchronously. A recent study from Li et al. [15] shows a promising approach that can be utilized to alleviate communication bottlenecks in the future FedDP infrastructure. Another method is to combine secured FL with a swarm optimizer based on the study of Zhan et al. [29] With these potential extensions, we anticipate that future FedDP can support a large number of various devices if the demand for bandwidth during the transmission can be satisfied with emerging technologies (e.g. 5G networks).

## Notes

### Competing Interest Statement

The authors have declared no competing interest.

