## Supplementary material for "FedDP: Secure Federated Learning for Disease Prediction with Imbalanced Genetic Data": sup

### Appendix

#### 1 Feature Selection

The gene expression data usually contain at least tens of thousands of genes for one sample, it should be noted that only a few groups of genes are related when it comes to some diseases. Moreover, training models based on these redundant genes not only consumes tremendous resources but also reduces the performance of disease prediction due to the overfitting problem. Thus, selecting the disease-mediated genes from the original gene expression data is required for the training process. In this experiment, we sorted genes based on fisher score, the fisher score is represent as:

$$F_{fisher}^{(k)} = \frac{\sum_{i=1}^{c} n_i(\mu_i^{(k)} - \mu^{(k)})^2}{\sum_{i=1}^{c} \sum_{x \in \omega_i} (x^{(k)} - \mu_i^{(k)})^2} \tag{1}$$

Where $x^{(k)}$ denotes the value of the sample $x$ on the $k$-th feature, $\mu_i^{(k)}$ represents the mean value of the $i$-th sample on the $k$-th feature, $\mu^{(k)}$ is the mean value of the samples of all categories on the $k$-th feature. The key idea of Fisher score is to find a subset of features, such that in the data space spanned by the selected features, the distances between data points in different classes are as large as possible, while the distances between data points in the same class are as small as possible.

However, the features selected by the fisher score also have drawbacks. On the one hand, since the fisher score calculates the score of each feature separately, it ignores the combination of features, that is, two or more features are evaluated together. For example, feature a and feature b may both have low fisher scores, but combination ab has a very high fisher score. In this case, the algorithm will discard features a and b, although in principle they should be chosen. On the other hand, they cannot handle redundant features. For example, feature a and feature b have very high fisher scores, but they are strongly correlated. In this case, the algorithm will select both features a and b, while feature a or b can be excluded with no loss in subsequent learning performance.

Therefore we further filtered the features based on their correlative coefficient matrix. Figure 1 (a) shows top-30 genes with highest fisher score. Their coefficient matrix is shown in Figure 1 (b). It can be seen that most features are highly related and redundant for classification. Hence, we filtered the sorted feature and set up a cut-off to extract 2000 features lower than the threshold. The appropriate threshold of the correlative coefficient for selection is -0.1. Figure 1 (c) shows the top 30 features we finally selected, they are apparently less correlated with each other.

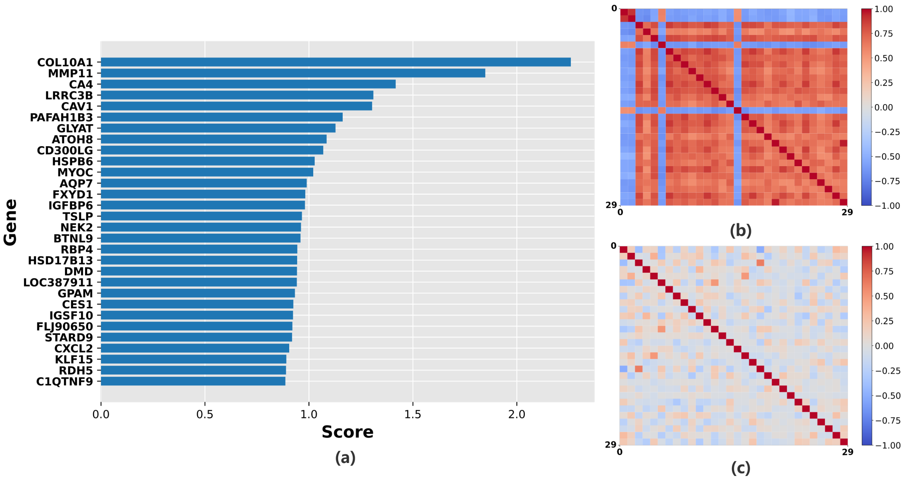


**Fig. 1.** Feature scoring and selection. (a). Top 30 genes with highest fisher score. (b). Heatmap of the correlative coefficient matrix of the top 30 genes after sorting by fisher score. (c). Heatmap of the correlative coefficient matrix of the top 30 finally selected genes.

#### 2 Training Processing

For federated model training, the gradients computed from local datasets are communicated to the central server at the end of each round. The pseudo codes of the proposed differentially private federated training procedure is depicted in Algorithm 1.

The proposed procedure protects per-sample privacy of each participating party dataset. Differential privacy is added by parties individually. In this way, the dataset sizes can be very different among the parties. Depending on the setting of sampling strategy, the sampling ratio q can be different from party to party. Thus the privacy loss is variant by the local dataset. Each party should keep record of the spent privacy along each update, and stops its update to central server once the predefined privacy threshold is reached.

#### 3 Cardiomyopathy Prediction on Balanced IQVIA Dataset

Even though TCGA program provides researcher a excellent platform for disease prediction, one issue is that the TCGA [1][3] datasets usually come with label-imbalanced problem. To build a generic and accurate framework, we also evaluated FedDP on IQVIA [2] dataset, which is sourced from IQVIA Inc. and de-identified for public access. The whole dataset contains 1713 samples, where 855 samples are diagnosed as wild-type amyloidogenic TTR cardiomyopathy

(ATTR-CM) (case group) and 858 samples (sample per row). Each sample has 1874 phenotypes (features per column). Compared with TCGA datasets, IQVIA dataset has balanced-labeled samples as well as different type of disease which enhance the generalty of FedDP.

**Table 1.** Performance of FedDP on IQVIA

| Noise multiplier | Delta | Batch size | Privacy eps | Model | Accuracy |
|---|---|---|---|---|---|
| 1.5 | 1e-3 | 32 | 1.73 | LR | 0.53 |
| | | | | CNN | 0.61 |
| 1.3 | 1e-3 | 32 | 2.14 | LR | 0.62 |
| | | | | CNN | 0.63 |
| 1.1 | 1e-3 | 64 | 3.19 | LR | 0.70 |
| | | | | CNN | 0.76 |
| 1.1 | 1e-3 | 32 | 2.97 | LR | 0.71 |
| | | | | CNN | 0.78 |
| 1.0 | 1e-3 | 32 | 3.11 | LR | 0.73 |
| | | | | CNN | 0.82 |
| 1.0 | 1e-3 | 32 | 3.38 | LR | 0.75 |
| | | | | CNN | 0.83 |
| 0 | 0 | 32 | None | LR | 0.80 |
| | | | | CNN | 0.89 |

**Results on IQVIA Dataset** Compared with two different TCGA datasets, the IQVIA dataset has more samples (1713 samples) with fewer features (1874 features). However, obtaining a good prediction on the IQVIA dataset is much more difficult than two TCGA datasets because features of IQVIA are more high-related. In order to simulate the real-world situations, we did not conduct any prepossessing for three datasets from the perspective of privacy-preserving at the local side. Thus, after fine-tuning, we can only reach the accuracy of 0.89 on our FedDP with the CNN model if there's zero-degree DP added, as shown in Table 1. For the best privacy budget we can get, the accuracy will decrease to 0.61. The best solution for considering both privacy-preserving and performance is to keep the multiplier to 1.1, delta is $10^{-3}$ and batch size is 32. This privacy setting achieved a 0.78 accuracy under a fair privacy budget that $\epsilon = 1.01$. The other LR model can only obtain an accuracy of 0.71 under the same privacy criteria.

**Algorithm 1** Federated training in FedDP

$\mathcal{D} = \{\mathcal{D}_1, ..., \mathcal{D}_K\}$: datasets held by Parties 1,...,$K$; $L$: Target loss function; $\Theta$: Trainable parameters; $\eta$: Round size
**procedure** FEDERATED TRAIN
  Initialize $\Theta$
  **for** $t \in \{1, ..., \eta\}$ **do**
    **for all** $\mathcal{D}_k$ **do**
      $\Delta_k^{(t)} \Leftarrow$ PartyUpdate($\Theta^{(t-1)}, \mathcal{D}_k$)
    **end for**
    $\Delta^{(t)} \Leftarrow \frac{1}{K} \sum_k \Delta_k^{(t)}$
    $\Theta^{(t)} \Leftarrow \Theta^{(t-1)} + \Delta^{(t)}$
  **end for**
**end procedure**
**function** PARTYUPDATE($\Theta, d$)
  $E$: Maximum allowed privacy cost; $M$: Noise multiplier; $S$: Batch size; $P$: Epoch size ;$\epsilon \Leftarrow$ compute_dp_sgd_privacy($d, B, M, T, E$)
  **if** $\epsilon \geq E$ **then**
    return 0
  **end if**
  Batch $\mathcal{B} \Leftarrow$ S samples from $d$
  Batches $\{\mathcal{B}_1, ..., \mathcal{B}_B\} \Leftarrow$ Random batches of $\mathcal{B}$
  **for** $b$ in 1, ...$B$ **do**
    g$\Leftarrow \nabla_\Theta \mathcal{L}(\Theta, \mathcal{B}_b)$
    $\Theta \Leftarrow \Theta - \eta$g
  **end for**
  $\Theta \Leftarrow \Theta$+Noise(M,S,d)
  $t \Rightarrow t + 1$
  return $\Theta$
**end function**

compute_dp_sgd_privacy(d,B,M,T,E): Function supported by Tensorflow_privacy [4] to compute specific privacy budget $\epsilon$ given a set of hyperparameters.

#### References


1. Bc-tcga dataset. http://dx.doi.org/10.17632/v3cc2p38hb.1file-c6b0b7d3-a63d-4ec7-87d5-ac2b81f38e00
2. Iqvia dataset. https://doi.org/10.1038/s41467-021-22876-9
3. Tcga-brca dataset. https://portal.gdc.cancer.gov/projects/TCGA-BRCA
4. Tensorflow privacy. https://github.com/tensorflow/privacy
